# RNA-binding protein A1CF modulates plasma triglyceride levels through posttranscriptional regulation of stress-induced VLDL secretion

**DOI:** 10.1101/397554

**Authors:** Jennie Lin, Donna M. Conlon, Xiao Wang, Eric Van Nostrand, Ines Rabano, YoSon Park, Alanna Strong, Behram Radmanesh, Yoseph Barash, Daniel J. Rader, Gene W. Yeo, Kiran Musunuru

## Abstract

**Background:** A recent human exome-chip study on plasma lipids identified a missense mutation in the *A1CF* (APOBEC1 complementation factor) gene that is associated with elevated triglyceride (TG) levels, but how A1CF, an RNA binding protein, influences plasma TG is unknown.

**Methods:** We generated *A1cf* knockout (*A1cf* ^−/−^) mice and knock-in mice homozygous for the TG-associated Gly398Ser mutation (*A1cf*^GS/GS^), determined lipid phenotypes, and assessed TG physiology through measurements of clearance and secretion. We further identified A1CF’s RNA binding targets using enhanced cross-linking and immunoprecipitation sequencing of cultured HepG2 cells and investigated pathways enriched for these targets. Transcriptomic effects of A1CF deficiency were evaluated through RNA sequencing and analyses for differential expression, alternative splicing, and RNA editing.

**Results:** Both *A1cf* ^−/−^and *A1cf*^*GS/GS*^ mice exhibited increased fasting plasma TG, establishing that the TG phenotype is due to A1CF loss of function. *In vivo* TG secretion and clearance studies revealed increased TG secretion without changes in clearance in *A1cf* ^−/−^mice. Increased VLDL-apoB secretion was also seen in *A1cf* ^−/−^rat hepatoma cells, but no increase in apoB synthesis was observed. This phenotype was seen without significant shifts in apoB-100/apoB-48 in A1CF deficiency. To discover novel pathways for A1CF’s role in TG metabolism, we identified A1CF’s RNA binding targets, which were enriched for pathways related to proteasomal catabolism and endoplasmic reticulum (ER) stress. Indeed, proteasomal inhibition led to increased cellular stress in *A1cf* ^−/−^cells, and higher expression of ER-stress protein GRP78 was observed in resting *A1cf* ^−/−^cells. RNA-seq of whole livers from wild-type and *A1cf* ^−/−^mice revealed that pro-inflammatory, not lipogenesis, genes were upregulated as a secondary effect of A1CF deficiency. Differential alternative splicing (AS) analysis and RNA editing analysis revealed that genes involved in cellular stress and metabolism underwent differential changes in A1CF deficiency, and top A1CF binding target proteins with relevance to intracellular stress were differentially expressed on the protein but not mRNA level, implicating multiple mechanisms by which A1CF influences TG secretion.

**Conclusions:** These data suggest an important role for A1CF in mediating VLDL-TG secretion through regulating intracellular stress.

## INTRODUCTION

Large-scale human genetics studies have identified DNA variants at novel genomic loci associated with cardiometabolic traits, including blood lipid levels ^1-7^. One of the strongest novel findings from a recent exome-sequencing study on lipid traits was an association between plasma triglycerides (TG) and the low-frequency variant rs41274050 (chr10: 52573772, hg19), a Gly398Ser (GS) missense mutation in the *A1CF* (APOBEC1 complementation factor) gene ^8^.

Although the A1CF protein has been studied in the context of apolipoprotein B (apoB) production, its precise function in TG metabolism still remains elusive. As an RNA binding protein (RBP) expressed in the liver, small intestines, and kidneys (Supplemental Figure 1A), A1CF was originally discovered in the context of binding *APOB* mRNA and complexing with the RNA editing enzyme APOBEC1 to facilitate editing of the *APOB* transcript. This interaction induces the deamination of cytidine^6666^ to uridine, thus converting the glutamine codon 2153 (CAA) to an in-frame stop codon (UAA) that generates apoB’s B-48 isoform (48% the length of the full B-100 isoform) ^9-14^. Previous reports have shown that loss of APOBEC1 function in mice does not affect circulating TG levels ^15,16^, suggesting that A1CF’s role in modulating plasma TG extends beyond its interaction with APOBEC1 and apoB-48 production.

In this study, we explored A1CF’s contribution to TG metabolism through an integrated unbiased discovery approach. We found in our genetic mouse models that the GS mutation results in a lipid phenotype consistent with A1CF loss of function, without shifts in B-100/B-48 ratios. To identify A1CF’s RNA binding targets outside of *A1CF* mRNA, we performed enhanced cross-linking and immunoprecipitation (eCLIP) followed by sequencing in HepG2 cells. Additionally, we performed unbiased transcriptome profiling to evaluate how A1CF interacts with mRNA transcripts to promote the TG phenotype. Through these profiling efforts we identified many RNA binding targets outside of *APOB* mRNA and also discovered a novel role for A1CF as an RNA splicing regulator. Ultimately, we found that through multiple mechanisms A1CF mediates intracellular stress and subsequent VLDL secretion.

## METHODS

Expanded Methods are available in the Online Data Supplement.

### Experimental animals

Homozygous *A1cf* knockout (*A1cf*^−/−^) and mice harboring the *A1cf* GS mutation (*A1cf*^GS/GS^) were generated using CRISPR-Cas9 genome editing, as described previously^8^. Briefly, to create both mouse strains simultaneously, we generated an *in vitro* transcribed CRISPR guide RNA (gRNA) targeting exon 9 of *A1cf*, which harbors the site of the Gly398Ser mutation. To knock in the Gly398Ser mutation, we also generated a 200-nucleotide single-stranded DNA oligonucleotide (ssODN) with homology to the mutation and other synonymous variants to prevent CRISPR-Cas9 re-cleavage of knock-in alleles. The Harvard Transgenic Core co-injected the gRNA and the ssODN with a Cas9-expressing mRNA into approximately 200 one-cell embryos of the C57BL/6J background (Jackson Laboratory). Following implantation of the embryos into surrogate mothers, approximately 50 mice were born, of which the majority harbored insertions and deletions (indels) while a few had Gly398Ser knock-in alleles (Figure 1). All mice were fed ad libitum with chow diet and fasted overnight prior to blood collection for plasma lipids, unless stated otherwise. Blood was collected through tail-vein puncture into heparin-lined tubes; blood samples were then centrifuged to obtain the plasma portion. Baseline plasma lipid and ALT levels were analyzed on an AXCEL autoanalyzer using commercially available reagents (Sigma Aldrich). Plasma total cholesterol (TC) and TG were also measured via calorimetric assay (Infinity reagents, Thermo Fisher) for samples collected during *in vivo* experiments.

**Figure 1.**
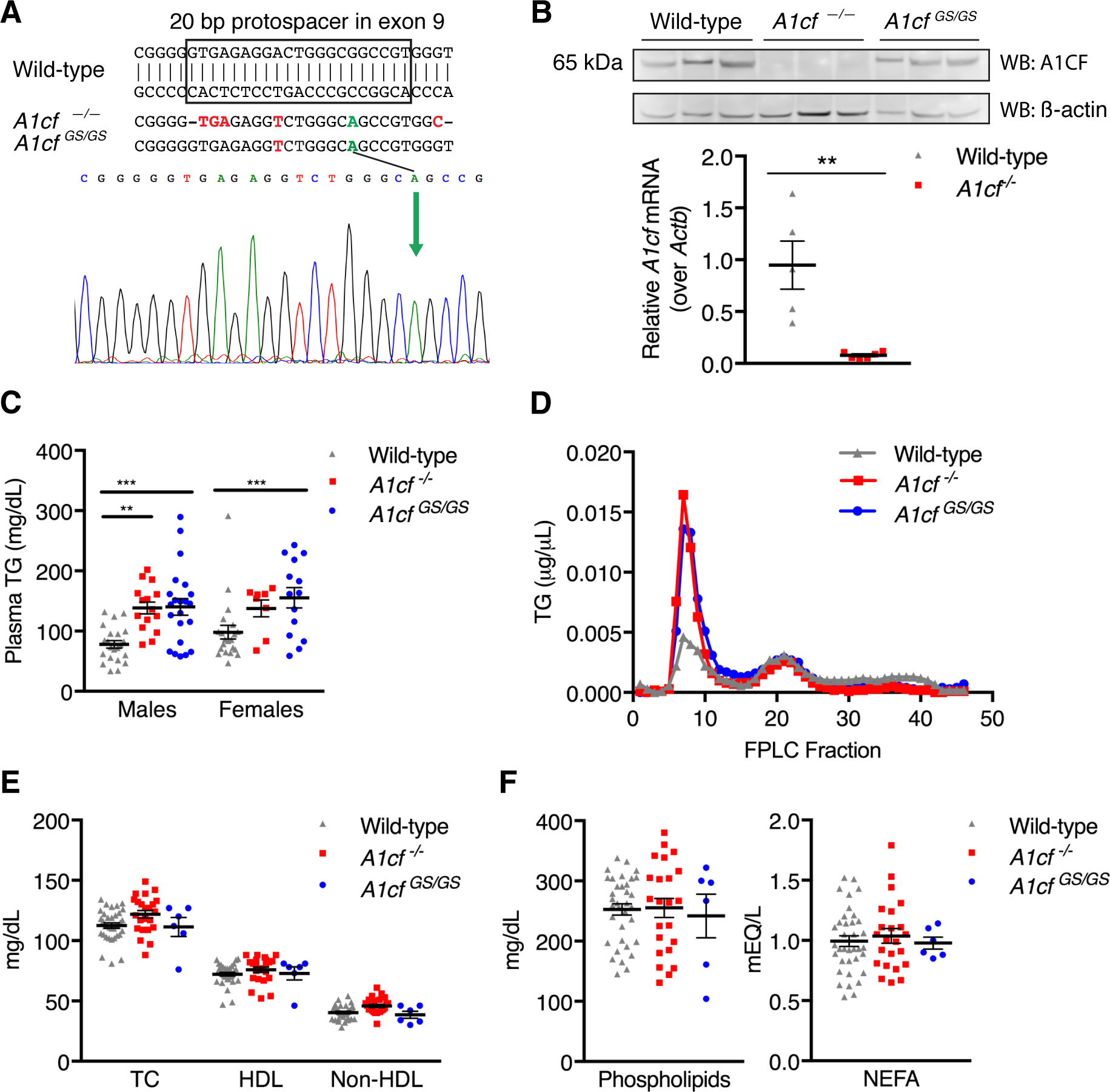
*A1cf*^−/−^and *A1cf*^GS/GS^ mice have elevated fasting plasma TG levels. (**A**) *A1cf*^−/−^and *A1cf*^GS/GS^ mice were generated using CRISPR-Cas9 genome editing. The protospacer and PAM within exon 9 of *A1cf* are demarcated for the wild-type sequence. Deletion of one bp induces a frameshift and premature stop codon (red) in one of the founder mice. Sanger sequencing confirms knock-in of the GS mutation (green). (**B**) Whole-liver protein lysates were separated by SDS-page and immunoblotted for A1CF. Representative blot shown includes *n* = 3 per group. RNA was also isolated from mouse livers and assayed for *A1cf* expression via real-time quantitative PCR, ** *P* < 0.01 (*n=* 6 mice per group). (**C**) Plasma TG of male and female wild-type (*n =* 21-22), *A1cf*^−/−^(*n =* 8-15), and *A1cf*^GS/GS^ mice (*n* = 14-22) were measured after an overnight fast. ** *P* < 0.01, *** *P* < 0.001. (**D**) Pooled plasma samples from mice of each genotype were fractionated by size, and TG content for each fraction was measured by fast protein liquid chromatography (FPLC). (**E**) Total cholesterol (TC), high-density lipoprotein (HDL), and non-HDL cholesterol levels were measured in wild-type (*n* = 36), *A1cf*^−/−^(*n =* 22), and *A1cf*^GS/GS^ (*n =* 6) mice after an overnight fast. (**F**) Phospholipids and non-esterified fatty acids (NEFA) were measured in wild-type (*n* = 36), *A1cf*^−/−^(*n =* 22), and *A1cf*^GS/GS^ (*n =* 6) mice after an overnight fast.

### Generation and culture of A1CF-deficient McArdle cells

McArdle RH-7777 (McA) rat hepatoma cells (ATCC) were maintained in Dulbecco’s Modified Eagle Medium (Gibco) with 10% fetal bovine serum, 10% horse serum, and 1% penicillin-streptomycin. The same guide RNA protospacer sequence used to generate *A1cf*^−/−^ and *A1cf*^GS/GS^mice was inserted into pGuide (Dr. George Church, http://www.addgene.org/64711/). The guide RNA plasmid was co-transfected with pCas9_GFP (http://www.addgene.org/44719/) into McA cells with Lipofectamine 3000 (Thermo Fisher) according to the manufacturer’s protocol. After 48 hours, cells were subjected to fluorescent activated cell sorting (FACSAria II) to isolate GFP-positive cells that were then plated at low density in 10 cm dishes to encourage formation of pure colonies from single-cell clones. Genomic DNA was extracted from colonies using the DNeasy Blood & Tissue Kit (Qiagen), and *A1cf* exon 9 was PCR amplified for Sanger sequencing to confirm the presence of indels. Clones with indels that shift the reading frame were selected for continued culture and cell experiments.

### eCLIP-seq of HepG2 cells

eCLIP was performed as previously described^17^. Briefly, A1CF-RNA interactions in 20 × 10^6^ HepG2 cells were stabilized with UV crosslinking (254 nm, 400 mg/cm^2^). Crosslinked cells were then lysed, with limited digestion of RNA with RNase I (Ambion). A1CF-RNA complexes underwent immunoprecipitation with an A1CF antibody (Abcam ab89050) using magnetic beads coupled with secondary antibody against Protein A. After stringent washes and dephosphorylation with FastAP (Thermo Fisher) and T4 PNK (NEB), a barcoded RNA adapter was ligated to the 3’ end on-bead with PEG8000. Samples were run on standard protein gels and transferred to nitrocellulose, with a region above the expected A1CF protein size isolated and digested with proteinase K. RNA from this isolation was then reverse-transcribed (AffinityScript, Agilent) and treated with ExoSAP-IT (Affymetrix) to remove excess oligonucleotides. A second DNA adapter was then ligated to the cDNA fragment 3’ end with high-concentration PEG8000 and DMSO. Each sample aliquot then underwent quantitative polymerase chain reaction (qPCR) and PCR amplification, followed by size selection by agarose gel electrophoresis. eCLIP-seq libraries were then sequenced on the Illumina HiSeq 2500.

eCLIP-seq data processing was performed according to a detailed pipeline with custom scripts previously described^17^. Peak-level input normalization was performed by processing non-A1CF control samples identically to A1CF eCLIP samples, and the number of overlapping reads between groups were counted and used to calculated fold enrichment normalized by total usable read counts. Enrichment P-values were determined by the Fisher Exact Test. Irreproducible discovery rate (IDR) analysis was performed by adapting the 2012 ENCODE IDR Pipeline documented at https://sites.google.com/site/anshulkundaje/projects/idr (Van Nostrand, *et al*. *in preparation*). eCLIP-seq data are available at https://www.ncbi.nlm.nih.gov/geo (accession number GSE117633). Sequences captured within IDR peaks underwent motif enrichment using MEME Suite ^18^.

### Study approval

All mouse care and use procedures were approved by Penn’s Institutional Animal Care and Use Committees.

## RESULTS

### *A1cf*^−/−^ and *A1cf*^GS/GS^mice have elevated fasting plasma TG levels

We generated viable *A1cf*^−/−^ and *A1cf*^GS/GS^mice using CRISPR-Cas9 genome editing (Figure 1A, Methods). We confirmed effective knockout of *A1cf* mRNA expression and A1CF protein expression in *A1cf*^−/−^ mice (Figure 1B) as well as successful introduction of the GS mutation in *A1cf*^GS/GS^mice (Figure 1A). After a 4-hour fast no significant differences in plasma TG were found (Supplemental Figure 1B). Strikingly, after an overnight fast *A1cf*^−/−^ and *A1cf*^GS/GS^mice of both sexes exhibited significant elevations in plasma TG compared to wild-type littermates (1.8-fold and 1.6-fold, respectively; Figure 1C), establishing that *A1cf*^GS/GS^mice recapitulate the exome-chip finding published by Liu *et al*. and that the GS mutation leads to at least partial loss of A1CF function in the lipid context.

Next, we separated lipoprotein fractions of plasma pooled from male mice of each genotype through fast protein liquid chromatography (FPLC) and observed that the elevations in TG content of *A1cf*^−/−^ and *A1cf*^GS/GS^plasma were found in the very low-density lipoprotein (VLDL) fractions (Figure 1D). No significant differences in TC, high-density lipoprotein (HDL) cholesterol, non-HDL cholesterol, phospholipids, and non-esterified fatty acids (NEFA) were present (Figures 1E and 1F). Although elevations in plasma TG often are accompanied by increased intrahepatic lipid content, no significant differences in intrahepatic TG or TC were found (Supplemental Figure 1C). After establishing that our *A1cf*^−/−^ mice phenocopy *A1cf*^GS/GS^mice, we focused our downstream studies on *A1cf*^−/−^ models.

### A1CF deficiency causes increased VLDL-TG secretion without increased apoB synthesis

To assess the physiological basis for the elevated TG seen in A1CF-deficient mice, we first evaluated whether fasting *A1cf*^−/−^ mice demonstrate increased hepatic TG secretion by measuring TG plasma accumulation after blocking lipolysis with intraperitoneal Pluronic P407. Compared to wild-type mice, *A1cf*^−/−^ mice exhibited 25% higher TG levels at 4 hours (*P <* 0.05) and 42% higher TG levels at 6 hours (*P <* 0.001) after Pluronic injection (Figure 2A), consistent with increased TG secretion from the liver in the setting of A1CF deficiency. To validate this further in cell culture, we generated *A1cf*^−/−^ McArdle-7777 (McA) rat hepatoma cells using CRISPR-Cas9 genome editing (Supplemental Figure 2A). We performed radiolabeling experiments with ^35^S-methionine to assess apoB-100 secretion and found that compared to wild-type, A1CF deficiency resulted in a 50% increase in apoB-100 secretion (*P <* 0.001) and a 30% increase in intracellular apoB-100 accumulation (*P <* 0.001) (Figure 2B). A similar pattern occurred with *A1cf* siRNA knockdown (Supplemental Figure 2B), indicating that increased secretion is unlikely due to off-target effects of genome editing.

**Figure 2.**
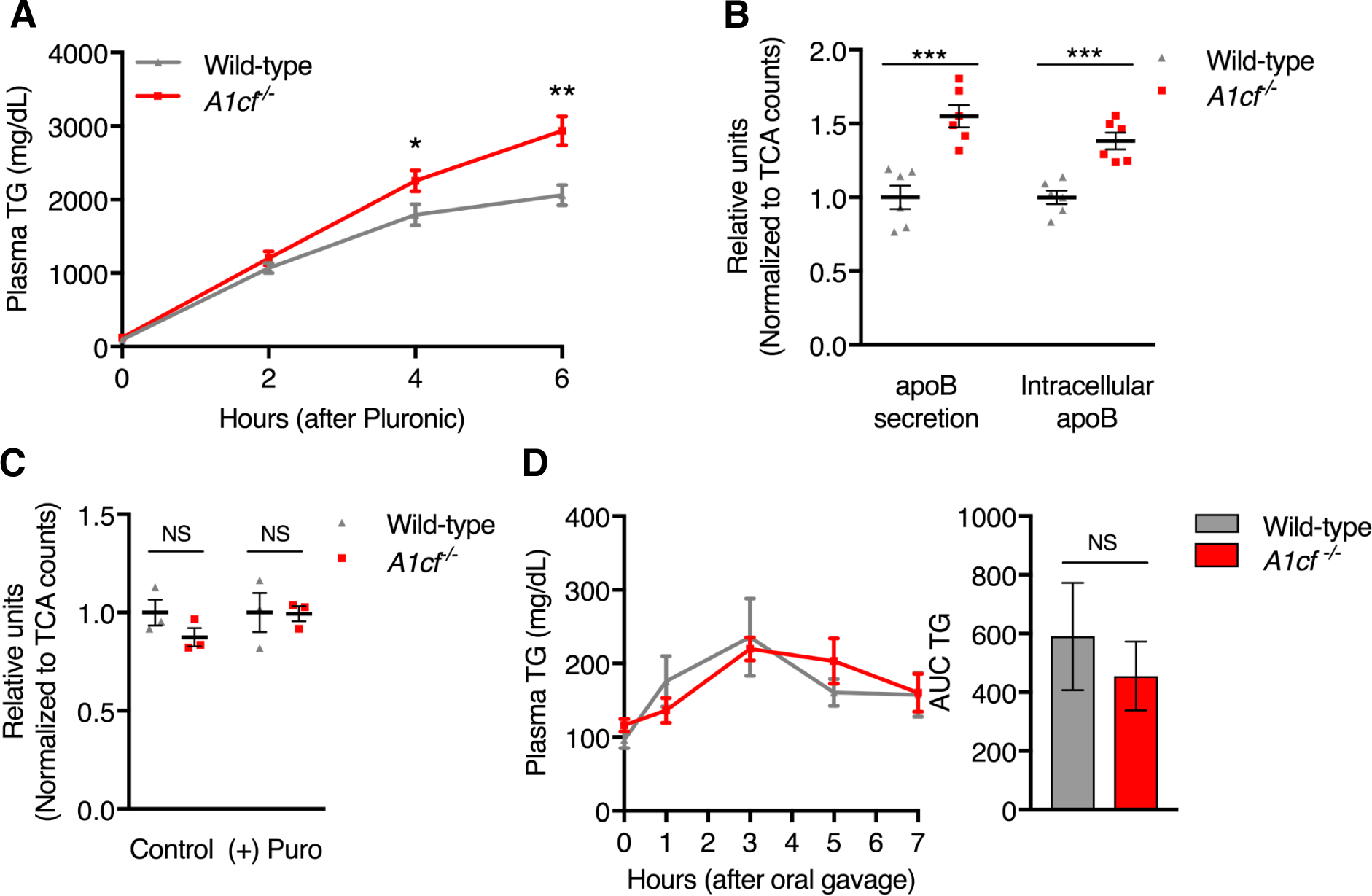
A1CF deficiency causes increased VLDL-TG secretion but does not decrease TG clearance. (**A**) Mice were injected i.p. with Pluronic P407 after an overnight fast. Plasma TG levels were measured after injection. *\* P <* 0.05, ** *P <* 0.01 (*n =* 4-5 mice per group). Results representative of 3 experiments. (**B**) *A1cf*^−/−^rat hepatoma McArdle (McA) cells were generated using CRISPR-Cas9 genome editing. Cultured knockout and wild-type McA cells underwent radiolabeling with ^35^S-methionine/cysteine, and newly synthesized cellular and secreted labeled apoB were measured. *** *P* < 0.001 (*n* = 6 wells per group). (**C**) McA cells were treated with and without 10 uM puromycin followed by pulse radiolabeling with ^35^S-methionine/cysteine to assess differences in apoB synthesis. Data represent relative amounts of intracellular radiolabeled apoB. (**D**) Mice were subjected to olive oil gavage after an overnight fast, and their plasma TG levels were measured after gavage. Clearance was assessed by AUC. (*n* = 4-5 mice per group). Results representative of 2 experiments.

To assess whether increased apoB secretion in A1CF deficiency was due to increased apoB production, we first compared intrahepatic *Apob* expression in *A1cf*^−/−^ and wild-type mice and found no significant differences in *Apob* transcript levels by qPCR (Supplemental Figure 2C). We then performed a pulse chase study with puromycin and found that apoB synthesis did not differ between wild-type and *A1cf*^−/−^ McA cells (Figure 2C), indicating that increased apoB production is not the driver of increased apoB secretion in A1CF deficiency.

### A1CF’s effect on plasma TG is not mediated through TG clearance

We next evaluated whether decreased clearance of TG-rich lipoproteins also contributes to the elevated plasma TG phenotype. Because A1CF is not expressed in adipose or skeletal muscle (Supplemental Figure 1A), we anticipated any clearance differences to be mediated through hepatic uptake. Wild-type and *A1cf*^−/−^ mice underwent an olive oil oral fat tolerance test, and their plasma TG levels were measured after oral gavage. The *A1cf*^−/−^ mice did not exhibit decreased TG clearance compared to wild-type controls, as indicated by similar TG curves and calculated areas under the curves (Figure 2D), further confirming that the primary physiological mechanism for elevated plasma TG in A1CF deficiency is due to increased hepatic VLDL-TG secretion.

### A1CF is not necessary for *Apob* mRNA editing and apoB-48 production

After establishing that A1CF modulates plasma TG levels through hepatic TG secretion, we then asked whether A1CF deficiency in mice leads to significantly reduced or absent *Apob* C^6666^ to U RNA editing to favor higher B-100/B-48 that would increase measured circulating TG mass. To capture the transcriptomic landscape, including RNA editing events, of A1CF deficiency, we performed RNA sequencing (RNA-seq) of poly-adenylated RNA isolated from whole livers of wild-type and *A1cf*^−/−^ mice (50-75 million 100 bp paired-end reads per sample, *n* = 6 mice per group). The liver was selected for RNA-seq due to its important role in lipoprotein metabolism. Using RDDPred (see Methods), for each RNA-seq library we quantified the proportion of *Apob* transcripts with C^6666^ to U editing. Notably, we found that 60% of transcripts from wild-type mice and 67.5% of transcripts from *A1cf*^−/−^ mice underwent this editing event (Figure 3A), indicating that the presence of A1CF is not necessary for this RNA editing event to occur. To confirm that apoB-48 is produced in *A1cf*^−/−^ mice without significant shifts in B-100/B-48, we performed Western blotting for apoB on plasma, whole-liver protein lysates, and small intestine protein lysates derived from wild-type and *A1cf*^−/−^ mice. Densitometry showed that no significant shifts in B-100/B-48 occurred in the absence of A1CF (Figures 3B and 3C). Although there appeared to be a trend towards increased intrahepatic B-100/B-48, this trend was not seen with circulating apoB, confirming that the plasma TG phenotype seen in A1CF deficiency is not due to changes in B-100/B-48.

**Figure 3.**
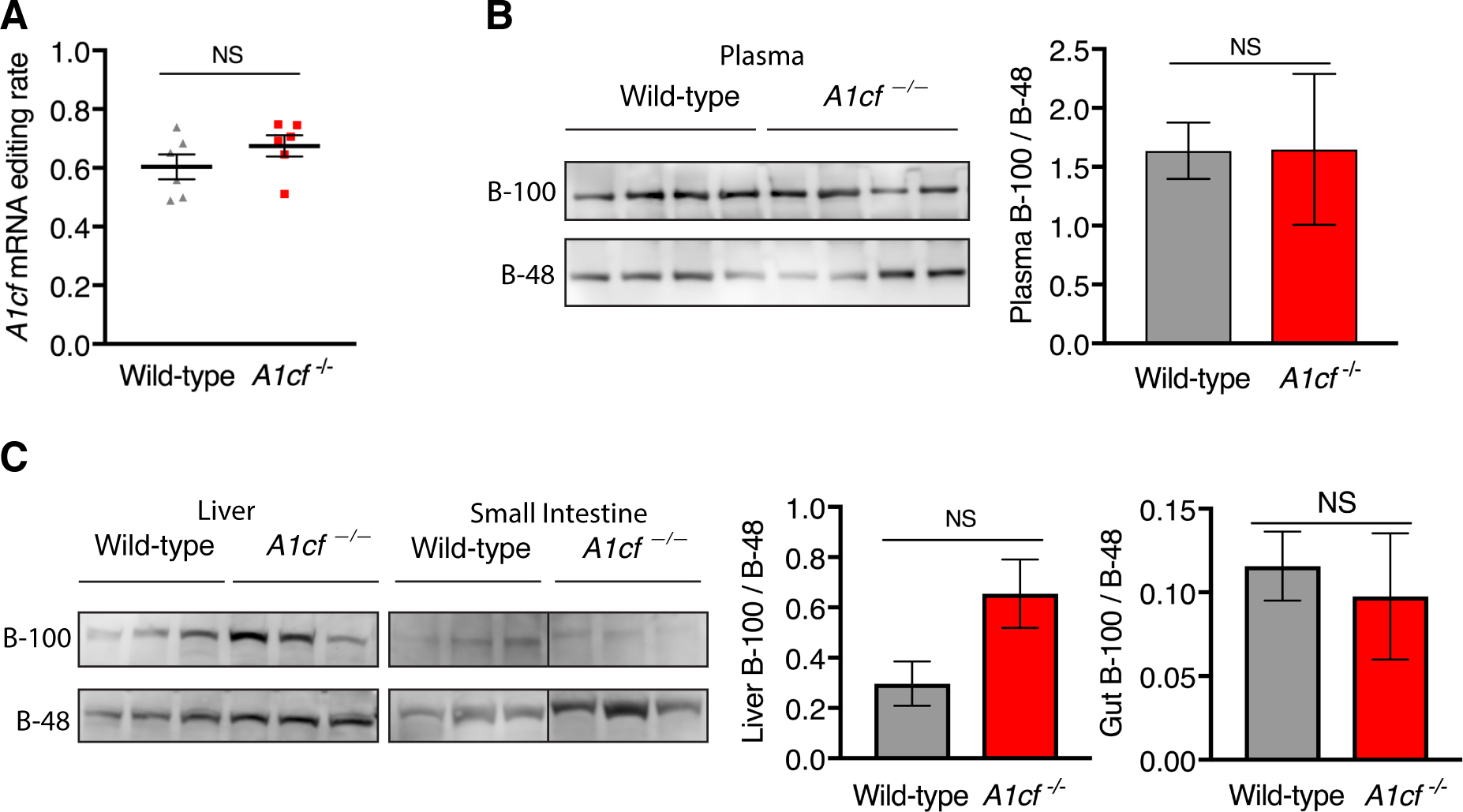
A1CF is not necessary for apoB-48 production through *Apob* mRNA editing. (**A**) RNA-editing rate for C^6666^ *Apob* mRNA of wild-type and *A1cf*^−/−^livers was calculated from RNA-seq using RDDPred (see Methods). *n* = 6 mice per group. (**B**) Plasma proteins from wild-type and *A1cf*^−/−^mice were separated by SDS-page and immunoblotted for apoB (*n* = 5 mice per group). Densitometry of the blot is reported as apoB-100/apoB-48 ratios and was analyzed with Student’s *t*-test. (**C**) Whole-liver and partial small-intestine protein lysates were separated by SDS-page and immunoblotted for apoB (*n =* 6 per group). Densitometry of the blot is reported as apoB-100/apoB-48 ratios and was analyzed with Student’s *t*-test.

### eCLIP-seq identifies non-*APOB* RNA binding targets enriched in protein processing and endoplasmic reiticulum (ER) stress pathways

Because increased apoB-100 secretion is not due to increased apoB-100 production or a shift in B-100/B-48, we next identified which non-*APOB* transcripts A1CF binds in order to investigate whether these binding targets form a network of genes and proteins relevant to apoB secretion. To identify A1CF’s direct binding targets, we performed eCLIP, a method by which A1CF was UV cross-linked to its binding sites and then pulled down by immunoprecipitation, followed by sequencing to map the cross-linked targets^17^. eCLIP was performed on HepG2 cells to identify binding targets in a human genomic context. Through eCLIP, 3617 significantly enriched peaks within 1233 unique binding target genes were identified for A1CF, with the majority of peaks falling within the 3’ untranslated region (UTR) of a transcript (Figure 4A and Supplemental Table 1). In addition, A1CF’s previously discovered binding motif by RNAcompete^19^ was validated in our experiment (Figure 4B). Gene ontology and pathway analysis for A1CF’s binding targets revealed that the top biological processes for these targets included proteasomal protein catabolism, viral immunity, cell-substrate adhesion, and response to ER stress (Figure 4C).

**Figure 4.**
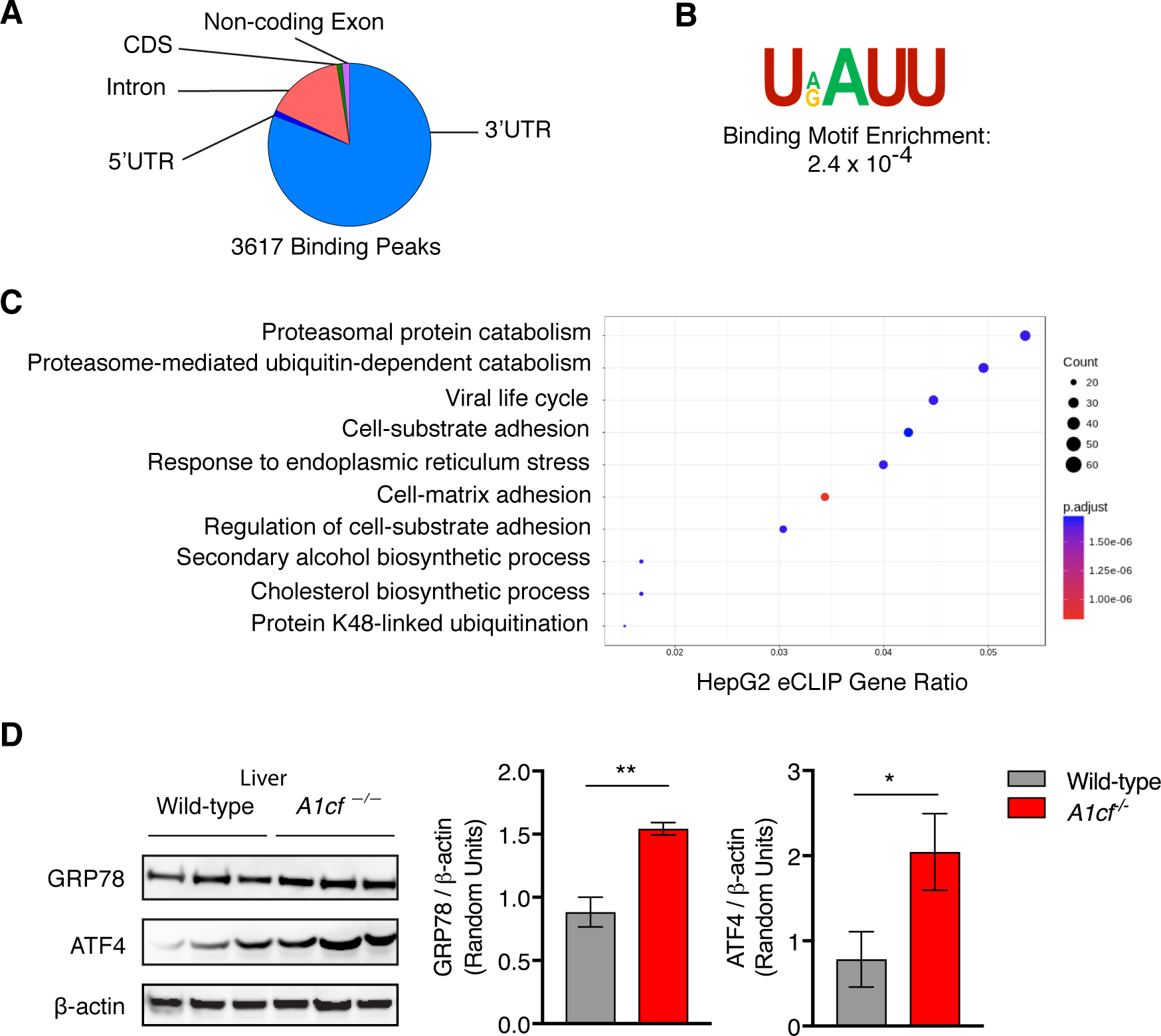
Enhanced CLIP-seq identifies A1CF’s binding targets. Enhanced CLIP-seq (eCLIP) was performed on HepG2 cells to identify A1CF’s RNA binding targets. (**A**) eCLIP identified 1411 binding clusters, and the distribution of these clusters among gene regions is represented in the chart. The vast majority of binding clusters fall within the 3’UTR of a gene. (**B**) Sequences from the binding clusters underwent motif enrichment analysis, which confirmed the predominant binding motif on eCLIP is consistent with the canonical binding motif. (**C**) Gene ontology analysis for genes represented in eCLIP binding clusters (eCLIP genes). The gene ratio reflects the relative percentage of eCLIP Genes involved in a given biological process. Top processes identified include those related to proteasomal catabolism and ER stress. (**D**) Whole-liver protein lysates were separated by SDS-page and immunoblotted for markers of ER stress, including GRP78 and ATF4. Blot representative of *n =* 6 per group. Densitometry of the blot was analyzed with Student’s *t*-test.

Because prior reports have demonstrated that moderate levels of ER stress increase VLDL secretion^20,21^, we next evaluated whether levels of ER stress differ in A1CF deficiency. By Western blotting analysis, *A1cf*^−/−^ mice have increased hepatic protein expression of ER stress markers GRP78 and ATF4 compared to wild-type mice (Figure 4D). Increased GRP78 expression was recapitulated in *A1cf*^−/−^ McA cells (Supplemental Figure 3A). Furthermore, exacerbating pre-existing ER stress by treating *A1cf*^−/−^ McA cells with the proteasomal inhibitor MG132 led to a decrease in apoB secretion and a significant drop in total protein levels compared to that seen in wild-type cells (Supplemental Figure 3B and Supplemental Figure 3C), results consistent with the parabolic curve of ER-stress induced apoB-100 secretion^21^.

### A1CF deficiency exerts secondary effects on mRNA expression

To assess whether A1CF influences intracellular stress pathways by affecting mRNA abundance of its binding targets, we examined the RNA-seq profiles of *A1cf*^−/−^ and wild-type mice for differential expression patterns. With a threshold of 1.5-fold difference in expression at FDR-adjusted *P* < 0.05, we identified 218 genes that were differentially expressed in hepatic A1CF deficiency (Supplemental Table 2). Surprisingly, very little overlap occurred between A1CF’s binding targets and genes differentially expressed in A1CF deficiency (Figure 5A), indicating that (1) A1CF’s effects on apoB-VLDL secretion are not mediated through transcriptional abundance of its binding targets and (2) A1CF’s influence on differential mRNA expression is a secondary rather than primary effect. This finding was validated through a complementary motif enrichment analysis^18^ demonstrating that the 3’ UTR sequences of differentially expressed genes were not enriched for A1CF binding sites (*P* = 1.0).

**Figure 5.**
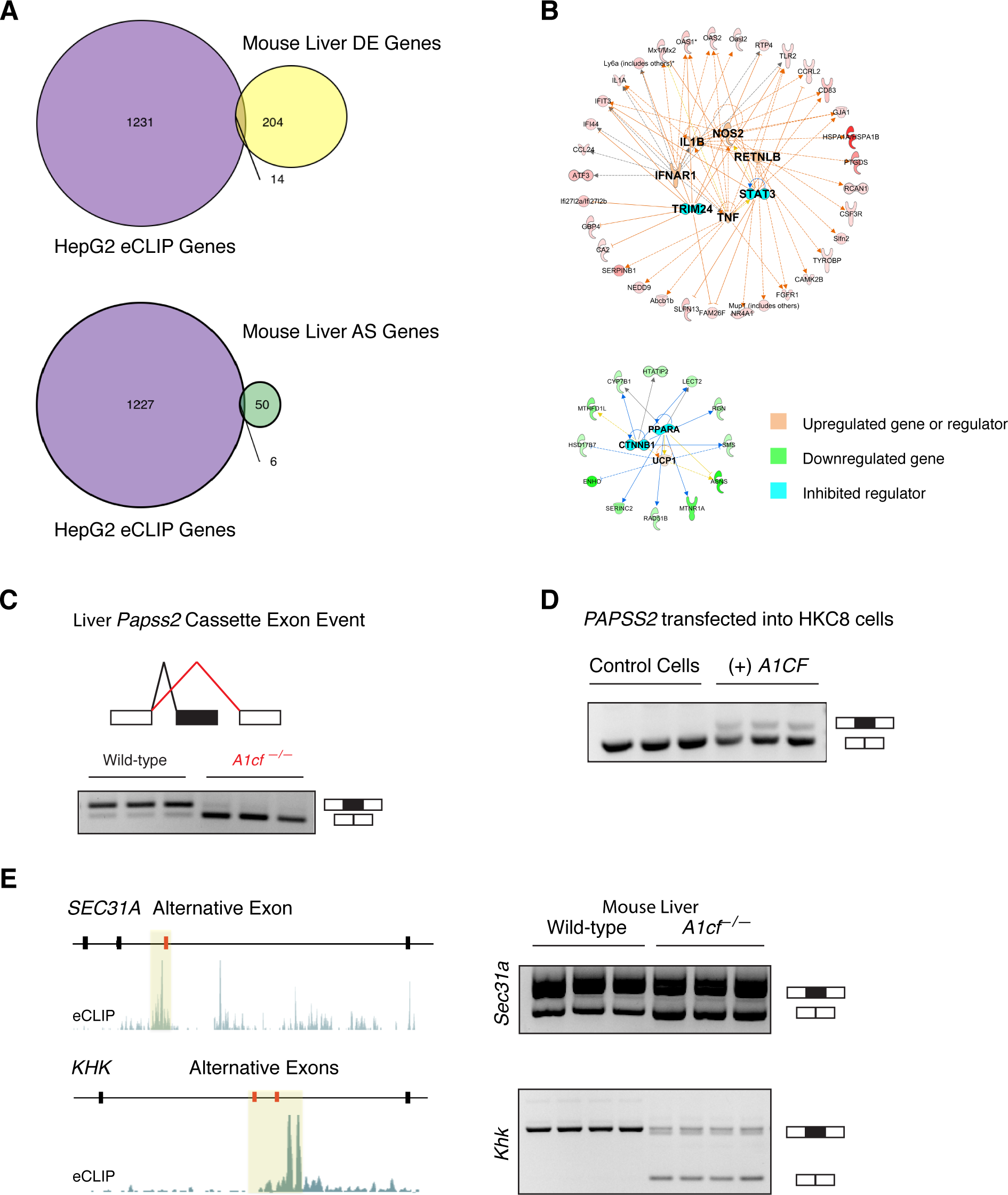
A1CF’s effects on the transcriptome. **(A)** Venn diagrams of overlap between eCLIP genes and genes differentially expressed in livers of wild-type vs. *A1cf* ^−/−^ mice (top) and between eCLIP genes and genes differentially alternatively spliced in livers of wild-type vs. *A1cf* ^−/−^ mice (bottom). (**B**) Upstream regulators of upregulated genes (top) and downregulated genes (bottom) are shown in their network relationship with their respective differentially expressed genes. (**C**) RNA-seq data generated for wild-type and *A1cf*^−/−^mouse livers were analyzed for differential alternative splicing (AS) events using MAJIQ. MAJIQ/VOILA output for differentially spliced gene *Papss2* is shown on the right, with delta percent spliced in (ΔPSI) at 86.2%. The black lines correspond to exon inclusion whereas the red lines correspond to exon exclusion seen in A1CF deficiency. The called splicing event was validated by RT-PCR of RNA derived from independent wild-type and *A1cf*^−/−^mouse livers that were not sequenced. (**D**)The same *PAPSS2* splicing event was induced in a human cell line (HKC8) that does not express *A1CF* through transfection with an *A1CF* expression vector; RT-PCR validation shown. (**E**) Out of the 6 genes that are both alternatively spliced in A1CF deficiency and eCLIP binding targets, only *SEC31A* and *KHK* demonstrated A1CF binding near the alternative exon. On the left for each gene, visualized eCLIP reads are presented in green, with the alternatively spliced exon(s) presented in red. On the right, RT-PCR validations are shown of homologous *Sec31a* and *Khk* differential AS events in wild-type and *A1cf*^−/−^mouse livers that were not sequenced.

Furthermore, genes more highly expressed in A1CF deficiency were linked to pro-inflammatory regulators such as IL1ß and TNF, while a subset of downregulated genes included targets of PPAR-α (Figure 5B). Consistent with the pro-inflammatory transcriptomic landscape of A1CF deficiency, protein dot blot assays on wild-type and *A1cf*^−/−^ liver lysates revealed relatively increased expression of pro-inflammatory cytokines and injury markers such as IL-27, NGAL, and Pentraxin-3 in *A1cf*^−/−^ livers (Supplemental Figure 4A). Lipogenesis genes were not significantly upregulated in RNA-seq and real-time qPCR validations (Supplemental Table 2, Supplemental Figure 4B), suggesting that increased *de novo* lipogenesis is likely not the primary pathway by which A1CF influences plasma TG levels and supporting the possibility that other mechanisms such as moderately increased ER stress may contribute to altered VLDL-apoB secretion.

### A1CF regulates RNA splicing of genes involved in protein processing and intracellular metabolism

Because A1CF deficiency does not significantly alter the differential mRNA expression of its binding targets, we next asked whether A1CF regulates the transcriptome by inducing changes in RNA splicing, an RBP-mediated process never previously attributed to A1CF. We analyzed our RNA-seq dataset for differential alternative splicing (AS) between wild-type and *A1cf*^−/−^ mouse livers using MAJIQ ^22^ and found 74 AS events occurring within 54 genes (Supplemental Table 3). We considered AS events as significant if the delta percent-spliced in (ΔPSI) for an alternative exon was greater than 10% for a predicted probability > 0.9. We prioritized AS events with ΔPSI > 20% for reverse transcription PCR (RT-PCR) using an independent set of mouse livers to validate that these splicing switches occur in the absence of A1CF. RT-PCR gels validate the AS events called for *Papss2*, *Sec31a,* and *Khk,* among others (Figures 5C and 5E). These genes encode proteins involved in sulfation, ER protein processing, and intracellular fructose metabolism, respectively ^23-26^. To further assess A1CF’s effect on splicing, we transfected human A1CF into the HKC8 cell line that does not express it constitutively and found that exogenous A1CF expression alone is able to induce the *Papss2* isoform seen in wild-type mouse liver (Figure 5D) even though *Papss2* mRNA is not a direct binding target of A1CF. In addition, we were able to confirm by eCLIP-seq that A1CF directly binds near the alternative exons for *SEC31A* and *KHK* in HepG2 cells (Figure 5E), validating that A1CF does play a regulatory role in alternative splicing for select transcripts relevant to intracellular stress and metabolism.

### A1CF regulates non-*Apob* mRNA editing of a limited number of genes involved in cellular stress

To identify other potential roles of A1CF as an RBP, we analyzed our RNA-seq dataset for differential RNA editing of genes other than *Apob.* To decrease the likelihood of false positives, we restricted our findings to editing events occurring in at least five out of six mice per group at a rate of greater than 25%. In addition, we considered only known and previously documented editing events. Consistent with our splicing analysis, our RNA editing analysis identified a relatively small gene set affected by A1CF deficiency. A total of nine transcripts were identified as undergoing A-to-I RNA editing in wild-type livers but not in the absence of A1CF (Supplemental Table 4), including the 3’ UTR of *Usp45*, a gene encoding a protein involved in ubiquitination and essential for DNA repair (Supplemental Figure 5) ^27^.

### A1CF mediates effects on intracellular stress by influencing protein translation

Although A1CF affects several differential mRNA expression and alternative splicing events in the liver, the transcripts affected by these events comprise only a small fraction of A1CF’s mRNA binding targets. To evaluate other possible effects of A1CF, we examined protein-level expression of several top binding targets by P-value. By Western blot analysis, GSR, CPE, and SCD1 were less abundant at the protein level in the setting of A1CF deficiency despite not being differentially expressed at the mRNA level (Supplemental Table 2, Figures 6 A-C). Each of these A1CF binding targets is involved in intracellular stress responses (see Discussion), suggesting that A1CF may influence translation of binding targets involved in the intracellular stress response that promotes increased VLDL-apoB secretion.

**Figure 6.**
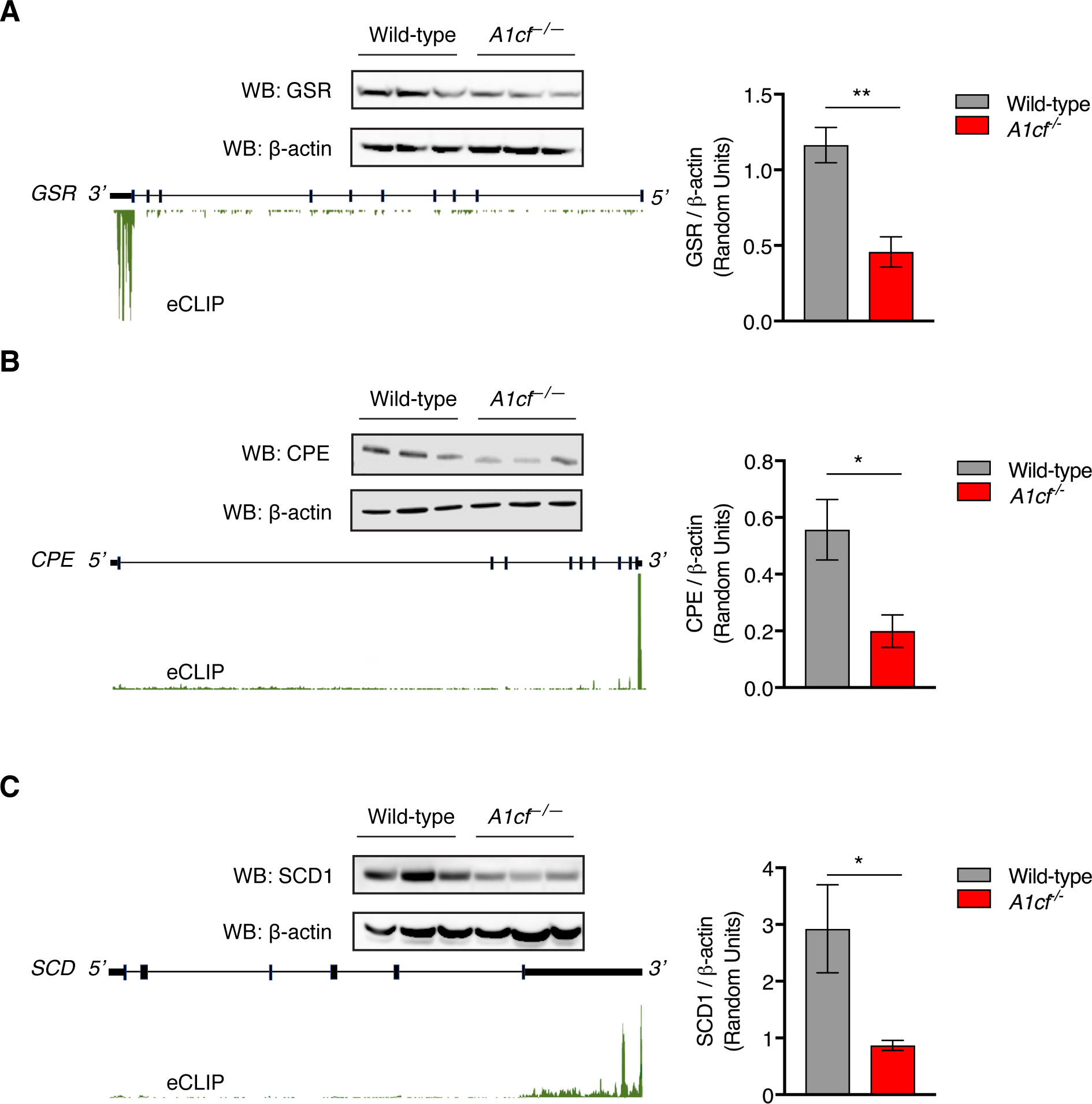
A1CF deficiency alters protein translation of binding targets. (**A-C**) Immunoblot of mouse livers was performed for A1CF binding targets *GSR*, *CPE*, and *SCD*. Representative blots are shown with corresponding densitometry (*n* = 3 per group). eCLIP binding sites for the target transcripts are visualized in green, falling in the 3’ UTR region.

## Discussion

Large-scale human genetics studies on plasma lipids have implicated many novel genomic loci in lipid metabolism. One of the strongest findings from a recently conducted exome-chip study was the association between *A1CF*’s low-frequency missense variant Gly398Ser (GS) and plasma TG levels. However, specific mechanisms through which such effect occurs remain unknown. Here we used CRISPR-generated mouse and cellular models to determine novel mechanisms by which A1CF influences plasma TG. By integrating transcriptomic studies with *in vivo* and *in vitro* functional studies, we (1) established that the TG-associated GS variant discovered in a large human genetics study is a loss-of-function mutation^8^, (2) discovered a new role for A1CF as a pathway-specific regulator of RNA splicing, (3) implicated A1CF in A-to-I editing that is independent of APOBEC1, (4) provided new insight that A1CF influences VLDL-TG secretion, and (5) identified A1CF’s involvement in ER stress as a possible mechanism for increased VLDL-TG secretion.

Because A1CF was initially discovered in the context of being an auxiliary protein in APOBEC1-mediated editing of *APOB* mRNA ^28^, most studies of A1CF function involved apoB isoforms in lipid metabolism, but little is known about A1CF’s role in modulating plasma TG. Fossat *et al*. previously showed that A1CF was not necessary for facilitating APOBEC1’s deamination of *APOB* mRNA C^6666^ to introduce the premature stop codon that leads to translation of the truncated apoB-48 isoform ^29^. A separate study has shown that *Apobec1* knockout mice do not exhibit a TG phenotype ^16^. These findings led us to investigate whether A1CF’s effects on modulating plasma TG are independent of its canonical role of facilitating *APOB* mRNA editing. Consistent with this, we demonstrated that B-100/B-48 ratios are similar in plasma, livers, and small intestines from wild-type and *A1cf*^−/−^ mice, suggesting that the elevated TG phenotype is present in *A1cf*^−/−^ mice regardless of apoB-48 abundance. Our mice differed from humans in expressing *Apobec1* in the liver and hence producing hepatic apoB-48 (humans apoB-48 is derived from the intestines only) ^30^. Nonetheless, we were able to recapitulate the human TG phenotype with the GS mutation in mice after an extended overnight fast, at which point all chylomicrons would have been cleared from systemic circulation. The near doubling of fasting plasma TG was seen in both male and female *A1cf*^−/−^ and *A1cf*^GS/GS^mice without a commensurate decrease in HDL-C, a result that focused our efforts outside of HDL biology.

To identify novel pathways relevant to A1CF’s modulation of plasma TG, we performed eCLIP-seq on HepG2 cells and RNA-seq on wild-type and *A1cf*^−/−^ mouse livers, because the liver is the most critical player in lipoprotein production and processing. Our decision to perform eCLIP-seq on HepG2 cells rather than on mouse liver was based on (1) our finding that an A1CF antibody that could undergo immunoprecipitation worked most cleanly in human-derived cells and (2) the concept that relevant TG-related A1CF biology would need to be conserved in both human and mouse in order to yield the similar TG phenotypes seen in the human genetics study and in our mice. In our eCLIP-seq analysis, we found that the majority of A1CF’s binding targets were not enriched for pathways related to TG metabolism or lipogenesis, but rather to proteasomal catabolism, cell adhesion, and ER stress. This result was consistent with our whole-liver RNA-seq analyses that revealed lipogenesis genes were not more highly expressed in *A1cf*^−/−^ livers. Additionally, *A1cf*^−/−^ and *A1cf*^GS/GS^mice showed no differences in circulating non-esterified fatty acids or intrahepatic lipid content, which is further evidence that increased lipid availability in the liver is not the driving force for the elevated TG phenotype ^31,32^. Instead, genes more highly expressed in A1CF deficiency were enriched in pro-inflammatory pathways and belonged to a network regulated by cytokines such as IL1ß and TNF, suggesting that A1CF deficiency results in intrahepatic stress and potentially ER stress, which has been closely linked to intracellular inflammation^33,34^. The enrichment of A1CF’s binding targets in pathways related to proteasomal catabolism and ER stress, as well as the observed differential protein abundance of ER stress markers GRP78 and ATF4 in A1CF deficiency, are of particular interest in the study of a TG phenotype because the ER is a key organelle in VLDL assembly and secretion^35^. Although the role of ER stress in VLDL secretion is complex, moderate levels of ER stress have been shown to result in increased VLDL secretion^20,21^. The effect of A1CF deficiency in promoting intracellular stress is therefore consistent with the observed effect on increased TG secretion *in vivo* and increased apoB secretion *in vitro*.

Our investigation of how A1CF interacts with RNA transcripts to influence intracellular stress and subsequent changes in VLDL secretion has shed light on novel roles of A1CF as an RBP in the cell. Initially we hypothesized that A1CF binds the 3’ UTRs of target transcripts that are then differentially expressed, as RBPs similar to A1CF in terms of sequence similarity have been found to stabilize mRNA transcripts by binding to 3’ UTRs ^36,37^. However, eCLIP-seq and binding motif enrichment analysis revealed that A1CF does not bind to the 3’UTR of the vast majority of genes differentially expressed in A1CF deficiency, indicating that these genes were differentially expressed as secondary or compensatory responses to A1CF deficiency. We then examined A1CF’s potential role in non-*Apob* RNA editing, alternative splicing, and protein translation. Although A1CF’s participation in the APOBEC1 RNA-editing complex has been well defined ^10,13,38-40^, it has never been described in the context of interacting with other RNA-editing enzymes such as the ADAR family. Here we show for the first time that A1CF is indeed involved in the editing of multiple transcripts other than *Apob* mRNA, including A-to-I RNA editing. The editing of *Usp45* is especially relevant to the TG phenotype given its role in ubiquitination ^27^, which can contribute to ER stress if dysregulated. Because this editing event occurs in a non-conserved portion of the 3’UTR, more studies are needed to examine A1CF-mediated RNA editing in human hepatocytes.

In addition to demonstrating A1CF’s involvement in non-*Apob* editing, we present novel data that A1CF contributes to the regulation of AS events. Although splicing regulators generally influence the AS of hundreds to thousands of genes ^41^, A1CF regulates the AS of a discreet set of genes that have relevance to essential intracellular processes such as protein transport and sulfation. Because splicing out alternative exons can change protein domains and protein folding ^42,43^, these AS events have the potential to change protein function. For example, the alternatively spliced exon in *KHK*, an A1CF binding target and differentially spliced gene in A1CF deficiency, results in altered fructose binding and hence fructose metabolism ^44^. Elucidating the causal mechanisms of how differential AS in A1CF deficiency contributes to ER-stress induced elevations in VLDL secretion will require further work.

Finally, we examined whether A1CF’s RNA binding targets had differential protein expression by performing Western blotting on top binding targets called by P-value in our eCLIP-seq data. Although the vast majority of these targets were not differentially expressed at the mRNA level, some of the top targets had different relative levels of expression at the protein level, indicating that A1CF binding somehow altered the translational process and thus efficiency. Future studies to examine protein abundance of A1CF’s binding targets on a high throughput scale will be needed to assess this finding further.

In summary, our studies in mouse and cellular models established that the *A1CF* GS variant associated with plasma TG is a loss-of-function mutation and perturbs TG metabolism through complex cascades of pathways. A1CF exerts its influence on plasma TG by regulating VLDL secretion in the setting of intracellular and, in particular, ER stress. As an RBP, A1CF is a novel regulator of the transcriptome that mediates intracellular stress through alternative splicing, RNA editing of genes involved in stress pathways and ER function, and altering protein translation of some of its targets. With its diverse functions in regulating the transcriptome, A1CF likely plays multiple roles in human lipid metabolism that will require further elucidation.

## Author Contributions

JL was responsible for conception, design, data acquisition, analysis, statistics, interpretation, and drafting and revising the manuscript. KM was responsible for conception, design, and revising the manuscript. DMC and XW were responsible for design, data acquisition, analysis, statistics, interpretation, and revising the manuscript. EVN, IR, and GWY were responsible for data acquisition, analysis, statistics, and revising the manuscript. AS was responsible for data acquisition and revising the manuscript. YB, BR, and YP were responsible for data analysis and statistics. DJR and SK were responsible for revising the manuscript.

## Acknowledgements

We acknowledge and thank Robert Bauer, Sumeet Khetarpal, and John Millar for helpful discussions.

## Sources of Funding

These studies were supported by NIH R01DK0999571 (to KM) and by NIH K08HL135348 (to JL). EVN was a Merck Fellow of the Damon Runyon Cancer Research Foundation (DRG-2172-13) and supported by an NIH K99/R00 Pathway to Independence Award (K99HG009530).

## Disclosures

GWY is a co-founder of Locana and Eclipse BioInnovations Inc. and member of the scientific advisory boards of Locana, Eclipse BioInnovations Inc. and Aquinnah Pharmaceuticals. EVN is a co-founder and consultant for of Eclipse BioInnovations Inc. The terms of these arrangements have been reviewed and approved by the University of California, San Diego in accordance with its conflict of interest policies. All other authors declare no competing interests.

